# Linkage disequilibrium - equivalence of the correlation and probability approaches

**DOI:** 10.64898/2026.09.24.754106

**Authors:** John Sved, Igor J. Chybicki

## Abstract

Amongst measures of linkage disequilibrium, the correlation measure *r*^2^, due originally to Sewall Wright, has several advantages. Although basically a haploid statistic, it can readily be estimated using covariance and correlation from diploid data. Its population expectation can also be found from probability methods. We show that the probability of linked identity by descent, LIBD, directly estimates the expectation for *r*^2^ for Wright-Fisher models. The simplest such model for two loci is the haploid model, where each haplotype is chosen independently from haplotypes of the previous generation. This gives an equilibrium between recombination (recombination frequency *c*) and drift (population size *N*) where the expected value of *r*^2^ is approximately 1*/*(1 + 4*Nc*). For diploid models, the expectation is similar for low values of *c*, but contains a term in *c*^2^ and rises to double the haploid value for the limiting value of *c* = 0.5. An increase in *r*^2^ is also expected for the model of monogamous mating, as opposed to random mating for each offspring. The expectation for *r*^2^ is unaffected by the use of diploid rather than haploid data, an exception being the case of monogamous mating. Computer simulation shows close agreement of *r*^2^ with probability expectations, with minor discrepancies observed for unlinked loci. Using the probabilistic definition, *r*^2^ can be intuitively understood as a parameter shaping the effective sample size in genome-wide association studies.

## 1 INTRODUCTION

Linkage disequilibrium (LD) can be measured in a variety of ways. A widely used statistic is the correlation coefficient *r*, or its square *r*^2^, recognising that LD is basically a correlation of frequencies in a population (see next section). This measure dates from Sewall Wright in 1933 [20], although its importance was not recognised until popularised by Hill and Robertson [7]. There have been several studies of the expected value of *r*^2^ in idealised (Wright-Fisher) populations [12, 10], an important study being that of Weir and Hill [19], where closed-form expressions for the diploid population expectations were derived together with finite-sample correction terms. Notably, except for [6] and [14], the *r*^2^ expectations were consistently derived using the ratio-of-expectations approximation rather than by directly taking the expectation of the squared correlation.

Independently, a simple probability method for calculating the expected value of *r* was suggested in [16]. The method relies on a comparison with inbreeding for a single locus. It is well known that the inbreeding coefficient *F* can be described either as a correlation between uniting gametes or as a probability of identity-by-descent [3]. It was suggested that a similar equation of correlation and probability methods can be applied for two loci. The argument is amplified in a following section.

The purpose of this paper is to show that the probability method leads to simple predictive equations for the correlation of frequencies in a variety of situations. Both haploid and diploid models are covered, with the latter accommodating gametic (phased) and zygotic (unphased) data types.

## 2 Correlation methods

### 2.1 Gametic (haploid) calculation

A 2-allele 2-locus model is assumed. It is convenient to describe correlations in terms of a table of *x* and *y* values for *A* and *B* loci respectively. Values of 1 and 0 are assigned for convenience, although this assignment does not change the correlation calculation.

Frequencies at the individual loci are

*p*_*A*_ = *g*_1_ + *g*_2_

*p*_*B*_ = *g*_1_ + *g*_3_

The usual LD coefficient, *D*, is defined as

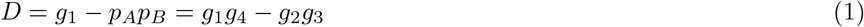

The covariance, Cov(*x*,y). is equal to

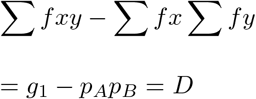

This is normalised to give the correlation:

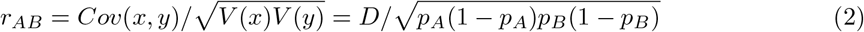

### 2.2 Zygotic (diploid) calculation

Gametic frequencies are usually not available, making it necessary to use a statistic that depends only on observation at the diploid level. Cockerham and Weir [2] introduced the concept of a ‘composite LD coefficient’, sometimes known as the ‘Burrows composite LD coefficient’ to take this into account. Gao et al. [4] showed that this coefficient is, in fact, the same as the zygotic covariance, allowing a calculation analogous to that in the gametic table. Note that correlation values are labelled *gametic* and *zygotic* below to distinguish them from *haploid* and *diploid* that are applied later to generation models

Table 2 shows the expected genotype frequencies in terms of haplotype frequencies *g*_1_ to *g*_4_ from the gametic table. Allowance is made here for departure from random mating according to the parameter *F*, which is applicable to either inbreeding or to population subdivision.

The zygotic covariance, is equal to ∑ *fxy −* ∑ *fx* ∑ *fy*, or ∑ *fxy − p*_*A*_*p*_*B*_, which simplifies to 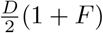.

The corresponding *V* (*x*) and *V* (*y*) simplify to 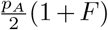 and 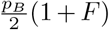 respectively.

Moving to the correlation, the factors (1 + *F*)*/*2 cancel, so that the zygotic correlation, which can be given the designation *R*, is equal to the gametic correlation *r*. This equality is an expectation only, given that frequencies will not be exactly as given in Table 2. Nevertheless there appears to be no bias in estimating gametic *r* from the diploid zygotic data, and we can write

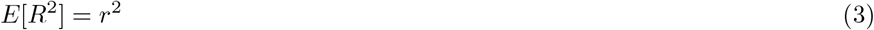

assuming the mating probabilities specified in Table 2. The factor of 2 in the estimation of *D*, however, indicates that the power of the estimation is only one half of what would be achieved if haploid gametic frequencies were available.

#### Probability methods

As mentioned previously, there is a close relationship between correlation and probability for single locus inbreeding models. Crow and Kimura [3] p67 argue that “If measurement can be thought of as being the sum of a number of elements, then the correlation coefficient is the measure of these elements that are common to the two measurements, the other elements being chosen at random”. In essence, in cases of IBD, with probability *f*, the correlation *r* = 1. In the absence of IBD, *r* = 0. Overall, *f ·* 1 + (1 *− f*) *·* 0 = *f*.

Sved and Feldman [16] suggested that this argument should hold for the equation of correlation and probability in a 2-locus model. Figure 1 shows a case where two haplotypes descend from a single haplotype, with no recombination on either of the 3 generation pathways. The probability that there is no recombination on pathway (1) is (1 *− c*)^3^, which may be designated as *f*. If there is no recombination in pathway (1), the correlation is 1. Any recombination event will recombine the *A* allele with a random *B* allele, giving a correlation of 0. Overall, therefore, the correlation *r* is *f ·* 1 + (1 *− f*) *·* 0 = *f*.

**Figure 1.**
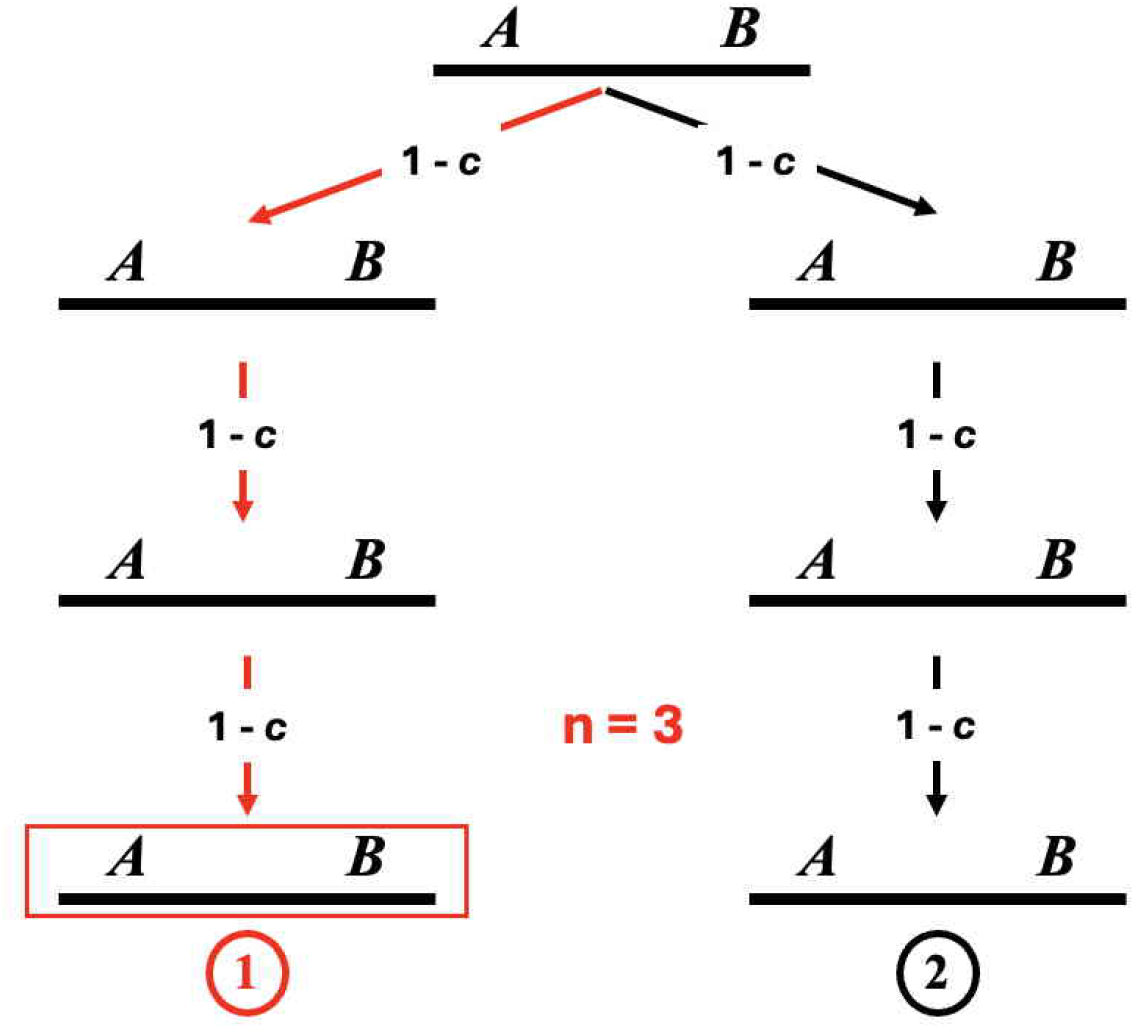
Identity by descent for a two locus model

Correlation coefficients are not, in general, additive. However in the present case the denominator of the coefficient, equation (2), is simply a function of individual allele frequencies at the two loci. These are not affected by recombination. Therefore additivity of the covariance term translates into additivity of the correlation.

Looking at the pair of haplotypes in the final generation of Figure 1 which may or may not be in the same individual, the probability of no recombination on either pathway is *f* ^2^. This event can be described as Linked Identity by Descent - LIBD [15], and given the designation *L*. In this case *L* = (1 *− c*)^6^ = *f* ^2^. The relationship *L* = *f* ^2^ is true for any number of generations.

Overall, therefore,

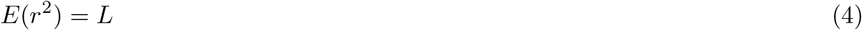

It is the ease of calculation of this LIBD probability for many population models that leads to simple expectations for *r*^2^. It is dependent on the assertion that that crossingover, if it occurs, will randomise connections between the two loci, leading to zero correlation.

## 4 Wright-Fisher 2-locus models

### 4.1 Haploid model

The simplest model is one in which each haplotype in the offspring generation is independently produced from a haplotype of the parent generation. Recombinants in such a haploid model result from crossingover between two randomly chosen haplotypes of the previous generation. The two parameters are *N*, the population size, and *c*, the crossover or recombination frequency. For comparison with the diploid model, the number of haplotypes is assumed to be 2*N*.

The probability that the same parental haplotype is chosen for each of two offspring haplotypes, equivalent to selfing in diploid models, is 1*/*2*N*. The LIBD probability in generation *t*+1, in terms of the LIBD probability in generation *t*, is then

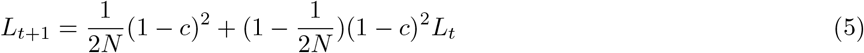

The (1*−c*)^2^ term here is the probability of no recombination in the production of either of the two gametes, leading to the two haplotypes of generation *t*+ 1. The first term in (5) may be termed ‘De novo LIBD’ and the second term ‘Retained LIBD’.

Using equation (4), the expected value of 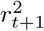 may be given as

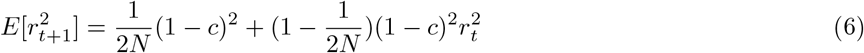

It should be noted that *r*^2^ is undefined if there is fixation at either of the two loci. No matter what the frequency at either locus in generation *t*, there is always a finite possibility of fixation in generation *t*+1. Strictly speaking, therefore, equation 6 makes sense only if interpreted as conditional on non-fixation. The transition from equation 5, to equation 6 interpreted in this way would be expected to introduce some bias, although presumably a small one.

A steady state 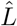 is reached when *L*_*t*+1_ = *L*_*t*_, which simplifies to

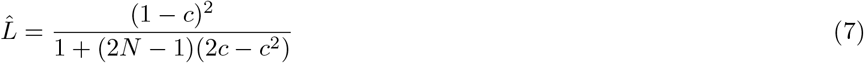

Using equation (4), the expected value of *r*^2^ at equilibrium may be given as

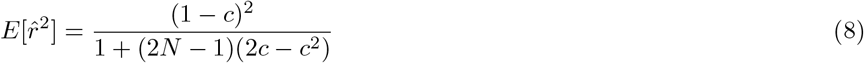

This relationship refers strictly to the population, from which a sample needs to be drawn to estimate any population statistics [8]. Such sampling is expected to increase the expectation. If the sample size is *S* indivduals, or 2*S* haplotypes, the expected increase to gametic *r*^2^ is 1*/*(2*S −* 1) [5]. It should also be noted that the original definition of this relationship in [16] confused the concepts of population and sample, essentially assuming that the population and sample sizes were the same.

For small values of *c*, equation (8) simplifies to

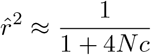

It may also be written exactly as

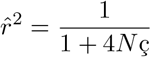

where

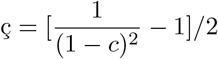

is a measure of recombination equal to *c* for low values and rising to 1.5 for the maximum value of *c* = 0.5

In any generation *t*, the value of *L* in terms of the original *L*_0_ is

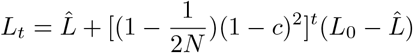

In terms of the expectation of *r*^2^ in any generation, this may be written as

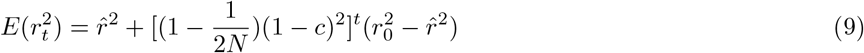

### 4.2 Diploid model

The key difference between haploid and diploid models is the pairing of haplotypes in the diploid model. Recombinants will involve the same pair of haplotypes in the diploid model, as opposed to random pairing of haplotypes to produce progeny gametes in the haploid model.

This pairing of haplotypes leads to a ramification not seen in the haploid model, as shown in Figure 2. The second panel here shows how two recombinant events in the parent can lead to identical haplotypes being passed down in the pedigree. It is still true for the diploid model that a single recombinant event will recombine the *A* allele with a random *B* allele, Only for “compensating” recombination events will this rule be broken.

**Figure 2.**
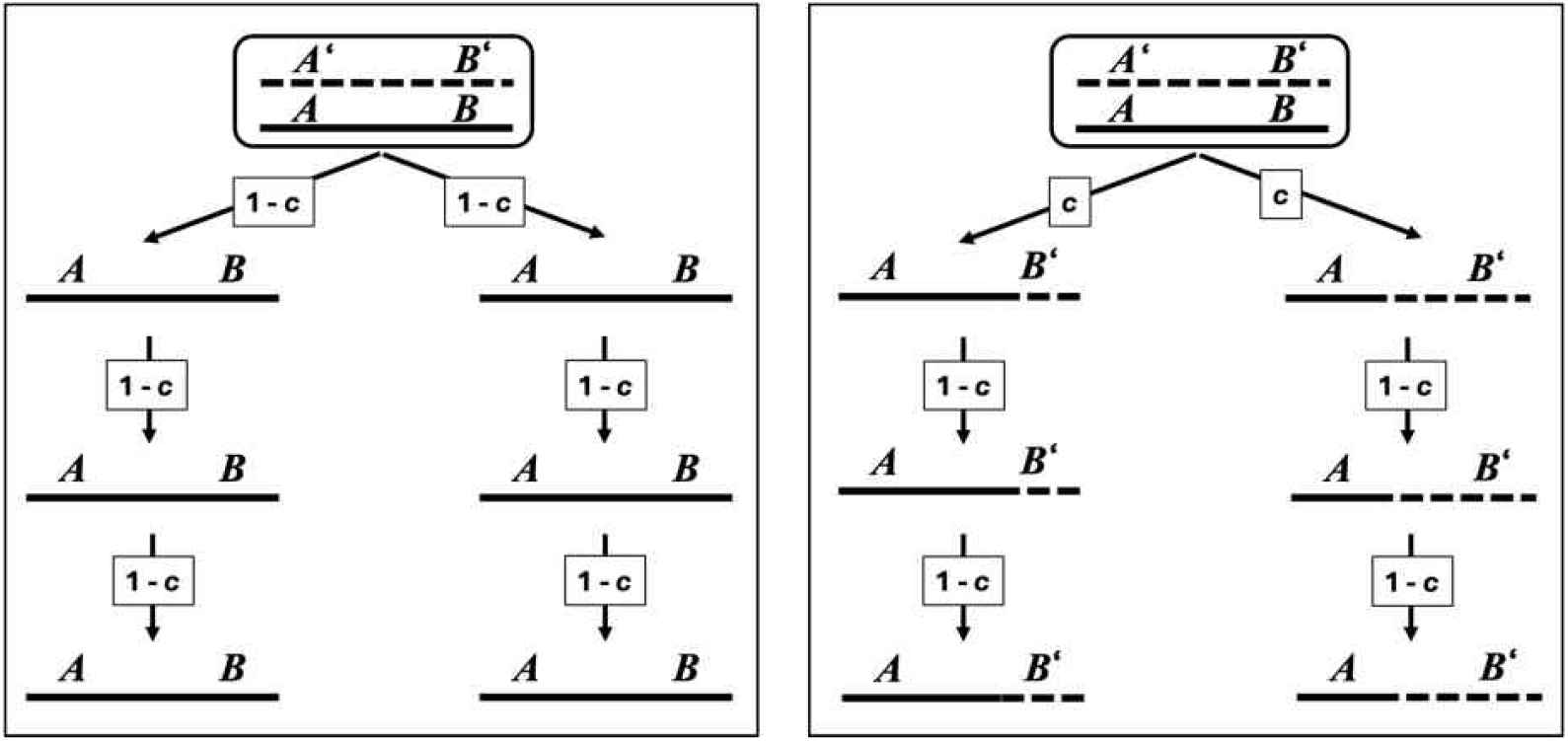
Two contributions to LIBD for the diploid model

Three different diploid models need to be distinguished. We consider here the model of separate sexes. The alternatives are single sex models, and within this category self fertilisation can either be allowed or forbidden. The allowance of self fertilisation is a key simplification. It makes such a diploid model comparable to the haploid model, because IBD is possible in a single generation.

With separate sexes, selfing is not possible. It means that IBD and LIBD cannot be defined in a simple pedigree with two parents and one offspring, as in the haploid case. Rather than looking at a single offspring, however, it is still possible to set up a 2-generation model with a population of parents and a population of offspring. IBD and LIBD cannot be defined in a single offspring, but can be defined in randomly selected pairs of haplotypes in different individuals in the offspring generation. In general in random mating populations, as assumed here, it is a matter of chance whether two identical haplotypes occur in a single individual or in randomly selected pairs from different individuals.

The calculations consider probabilities for a single offspring pair. These offspring may have zero, one or two parents in common. The most likely case is that there will be zero parents in common. If there are *N*_*f*_ females and *N*_*m*_ males in the parent generation, the chance of no parents in common will be 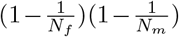 .The chance of two parents in common, ie full sibs, is 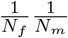 The remaining probability, one parent in common, half sibs, is 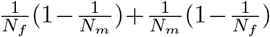.

We look initially at probabilities for these three cases separately, starting with two parents in common. Note that in these calculations the sexes of the two offspring individuals are not relevant, assuming that sex is inherited independently from the loci under consideration.

1. Two parents in common (LIBD probability is defined as *L*[2]) Calculations are shown in Appendix A. These lead to:

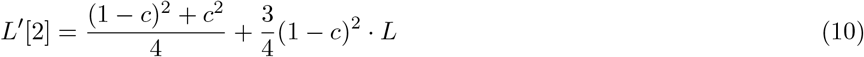

where L’(2) is the probability of LIBD of a randomly selected haplotype from each offspring and *L* is the LIBD probability in the parent generation.
2. One parent in common (Appendix B):

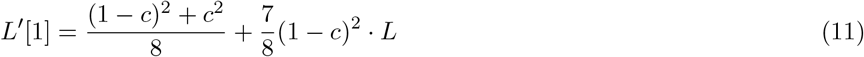
3. No parents in common.

There is no possibility of LIBD newly arising in the mating of two unrelated offspring individuals. The LIBD probability in this case becomes simply the probability in the previous generation less that lost through recombination:

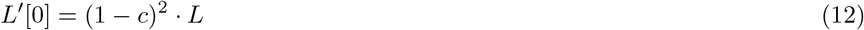

#### 4.2.1 Overall recurrence relationship

Table 3 summarises the probabilities of shared parents and the LIBD probabilities in each case. Weighting contributions to *L*^*′*^ by their frequencies over the four classes in the table, the overall recurrence relationship simplifies to

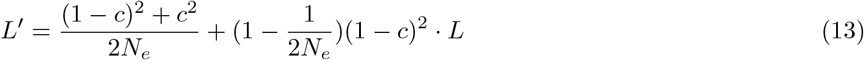

where *N*_*e*_ is the usual effective population size given by

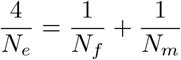

The equilibrium value of *L* is

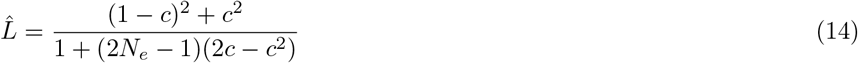

Both of these differ from the haploid formulae only by the *c*^2^ term, which becomes significant only for loose linkage.

The calculations of Appendix A show explicitly which recombinant products contribute to genotypes with the two haplotypes identical at both loci. They confirm the arguments related to Figure 2 that only “compensating recombination events” lead to “De novo LIBD”. Recombinant events other than these can be ignored, since they lead to zero correlation as outlined in the previously quoted statement [3] “. the other elements being chosen at random”.

### 4.3 Single sex diploid model

Expectations under this model are identical to those from the separate sexes model, equation 13 for the recurrence relationship and 14 for the equilibrium. Allowance for self fertilisation considerably simplifies the calculations. Although detailed calculations are not shown for this case, Appendix C outlines the general argument, in particular showing the balance between production of de novo LIBD against the loss of LIBD due to recombination.

### 4.4 Simulation of the diploid model

Two different simulation programs have been used. The first program (see Appendix D) simulates a single continuous chromosome, and is used to test agreement of simulated *r*^2^ values with expectation based on equation 13 for a range of low *Nc* values. Figure 3 shows the close agreement of observed and expected values for this case starting with complete Linkage disequilibrium (*r*^2^ = 1.

**Figure 3.**
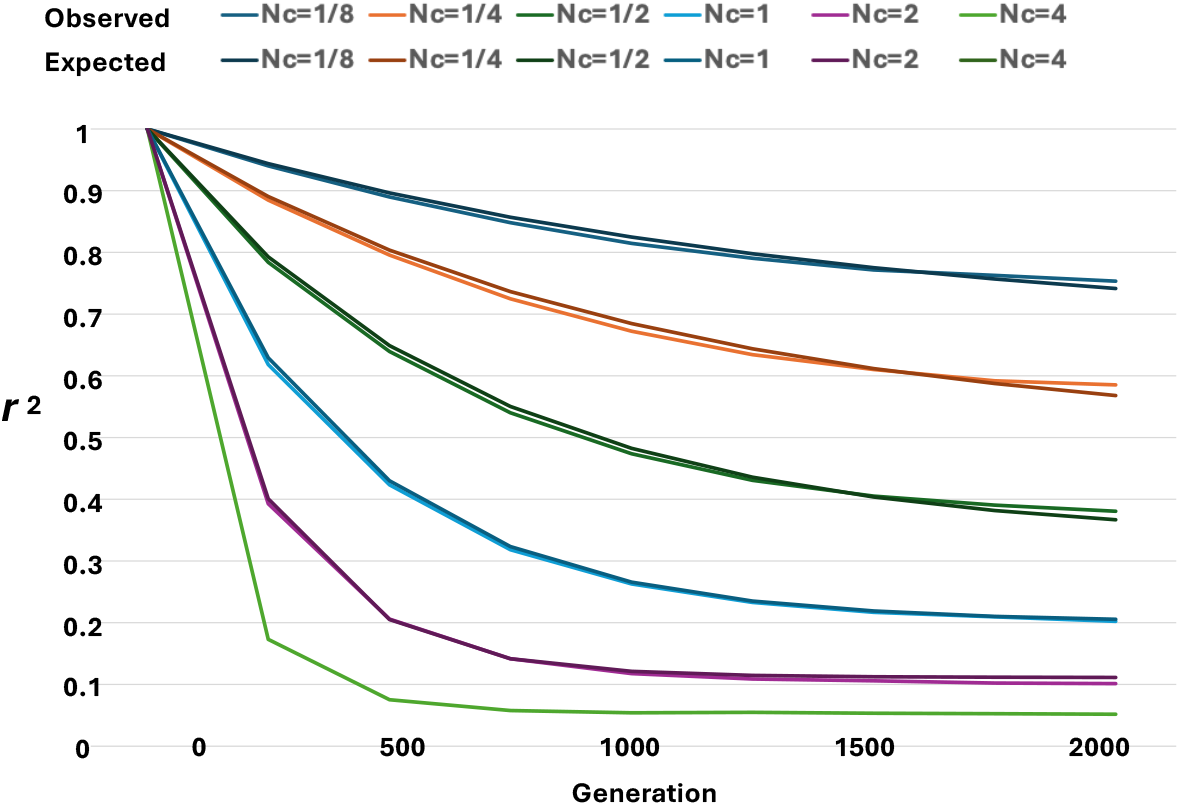
Observed *r*^2^ for populations of size *N* = 1000 starting with complete LD, compared to prediction based on (9

Figure 4 shows the comparison of observed and expected *r*^2^ values for simulations of the highest value of *c* = 1/2, starting from Linkage Equilibrium. This level of recombination accentuates the difference between haploid and diploid models.

**Figure 4.**
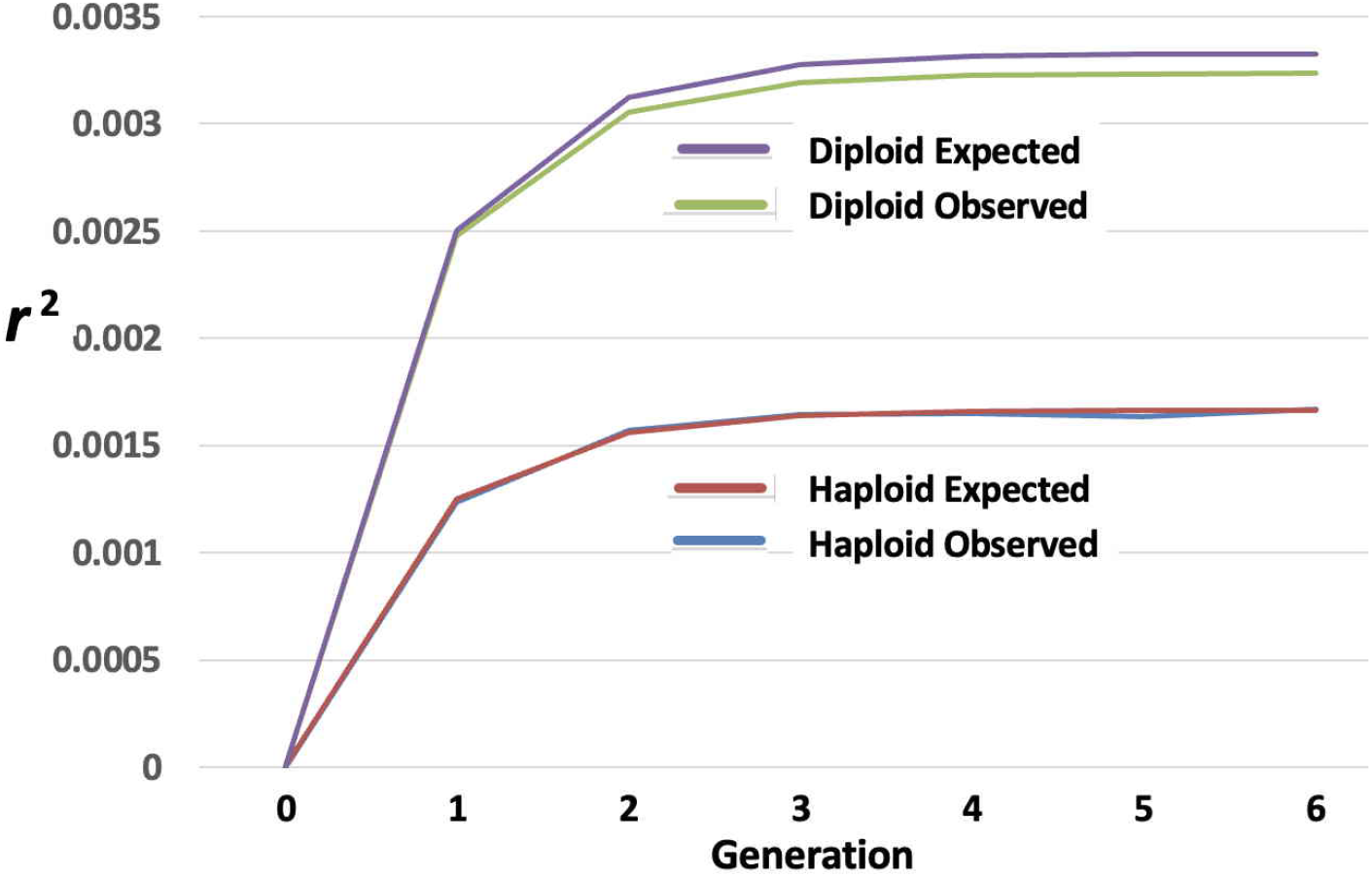
Comparison of simulated and expected *r*^2^ for haploid and diploid models with *N* = 100, *c* 0.5.

Figure 4 is based on the ‘2loc sim’ program (see Appendix D). Although there is reasonably good agreement, observed *r*^2^ values for the diploid model are slightly below expectation. As expected, there is approximately a 2*×* equilibrium difference for this limiting value of *c* = 1/2.

Observed *r*^2^ values shown in Figure 4 have been corrected by subtracting the expected sampling variation from the calculated *r*^2^. In this case, where the entire population has been sampled, the expected sampling variance is 1*/*(2*N −* 1) = 1/199 for gametic *r*^2^ and 1*/*(*N −* 1) = 1/99 for zygotic *r*^2^. Each of these corrections considerably outweighs the resultant *r*^2^, whose expected values are close to 1/300 for the diploid model and 1/600 for the haploid model.

### 4.5 Monogamous mating

A surprising finding was reported [19] for monogamous mating, where each female mates over a lifetime with a single male, rather than the random pairing for each offspring implied by Table 3. It was shown that such mating may considerably increase the expected level of LD.

The LIBD calculations for monogamous mating can be summarised by modifying the probabilities of Table 3 in Table 4. Monogamous mating implies that *N*_*f*_ = *N*_*m*_ = *N*_*e*_*/*2. The probability that a random pair of offspring gametes has two common parents is then 2*/N*_*e*_. The probability of a single common parent is zero, and the probability of no common parents is 1 *−* 2*/N*_*e*_. Weighting equations (10) and (12) by these values gives the offspring LIBD probability for monogamous mating as

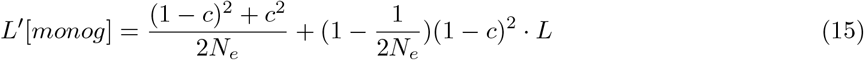

This equation is identical to equation (13) for non-monogamous mating. This is inconsistent with the result given by Weir and Hill [19]. It is also inconsistent with computer simulation (see following section), showing a rise in *r*^2^ values for monogamous mating as compared to non-monogamous. It implies that there is another factor that equation (15) does not take into account.

This quandary is resolved by the recognition that monogamous mating introduces a new class of mating that contributes to LIBD through recombination. It results from the possibility that two chosen offspring have no common parents, but instead have related parents that are themselves the product of sib mating.

The three generation pedigree of Figure 5, specifically the parents and inside grandparents shown in black, show the passage of alleles. Specific alleles from the grandparents, *A*1*B*3 are chosen for the parents. Recombination in these parents results in identity of two haplotypes in the offspring.

**Figure 5.**
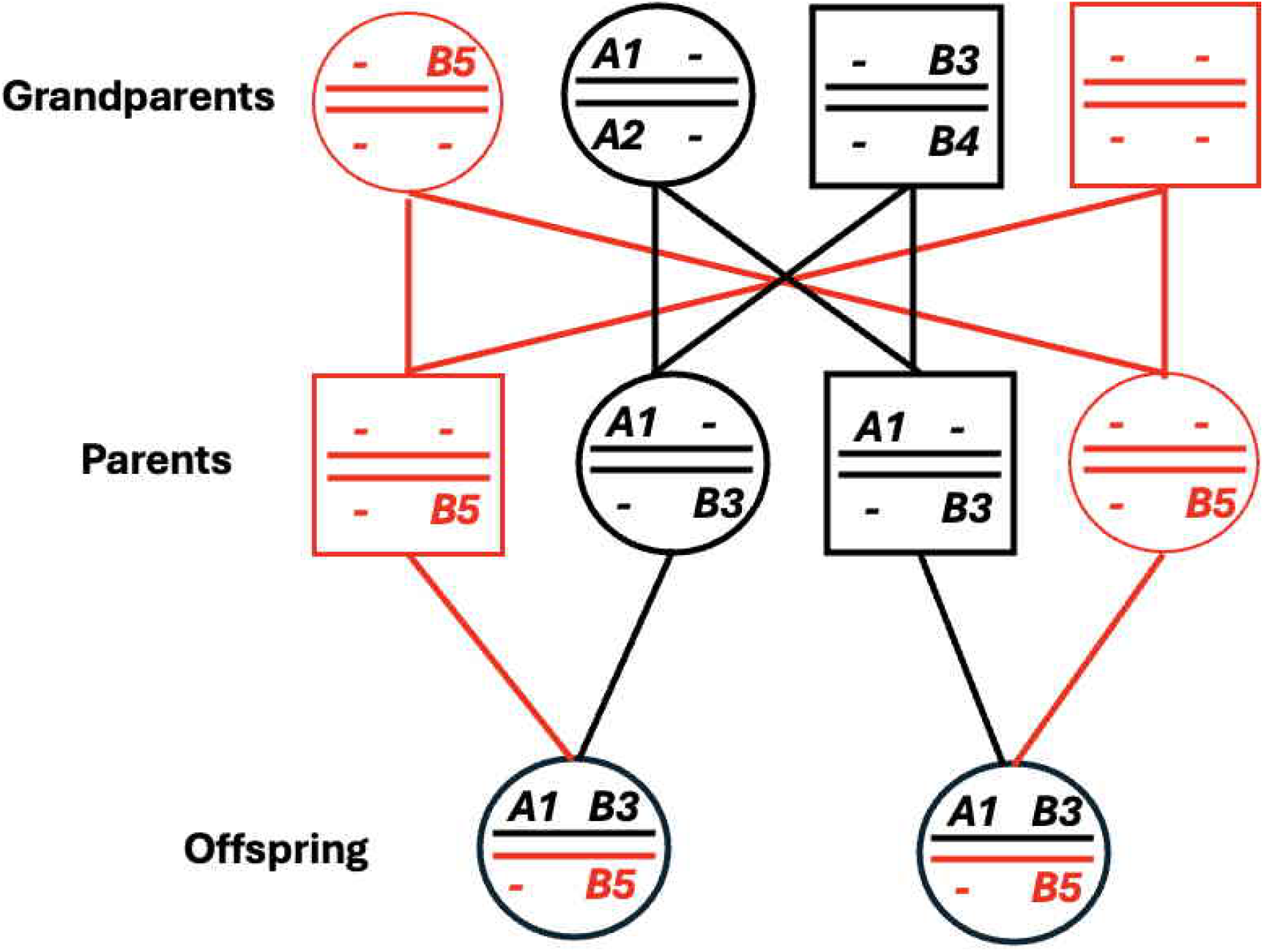
Pedigree for the study of monogamous mating.

The probability that *A*1 is found in both parents is 1/4, irrespective of any recombination event in the grandparents. Similarly for *B*3. The probability of both events is 1/16. Then the probability of recombination and passage of the *A*1*B*3 gametes to both offspring is *c*^2^*/*4. But there are 4 possible haplotypes which will have the same effect, *A*1*B*3, *A*1*B*4, *A*2*B*3, *A*2*B*4. Furthermore there are 4 equivalent haplotypes where the *A* allele comes from Parent 2 and the *B* allele from Parent 1. The overall probability of this LIBD event is thus 1*/*16 *× c*^2^*/*4 *×* 8 = *c*^2^*/*8.

The calculation then needs to take into account the probability that the selected parents are themselves from a single mating, which is 2*/N*_*e*_ as previously. Overall, therefore, the contribution given by related parents is

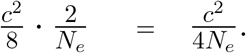

The overall recurrence relationship for *L*, as given by (13) for non-monogamous mating needs to be modified for monogamous mating by adding the *c*^2^*/*4*N*_*e*_ contribution, giving

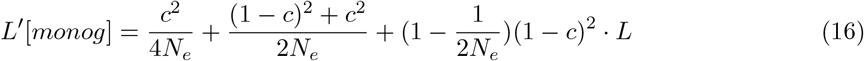

The equilibrium value is

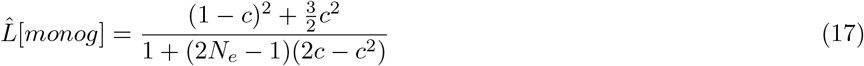

in close agreement with the value given in [19], equation 6.

### 4.6 Zygotic correlation

It is interesting to study the zygotic correlation for monogamous mating. The expectation from equation 3 is that the zygotic correlation has the same expectation as the gametic correlation, even allowing for departures from random mating such as inbreeding or population subdivision. However the case of monogamous mating appears to provide an exception to this rule.

Figure 5 shows, in red, an additional effect of monogamous mating. The principle of the composite index [2] relates to ‘haplotypes’ such as *A*1*B*5 in Figure 5. The LIBD principle can also be applied to the identity of such ‘haplotypes’, as shown in the figure. The probability of IBD of alleles such as *B*5 is 1/8, since the IBD could come from either grandparent. But these two parents also need to be sibs, with probability 2/*N*_*e*_. The overall probability is thus 1*/*8 *×* 2*/N*_*e*_ = 1/4*N*_*e*_, This probability supplements the value for *L*^*′*^ from equation 16, giving

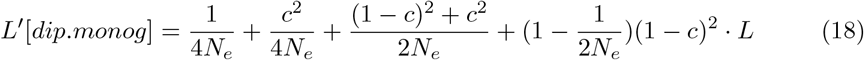

Figure 6 shows simulations, based on 10^8^ replicates of single-sex diploid populations of size 100 with *c* = 0.5. The two lowest lines show non-monogamous mating for comparison. The observed value with gametic *r*^2^ is the same as shown in Figure 4. Both sets of values have been corrected for variation due to sampling. For zygotic *r*^2^, the correction is 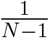. The agreement of the values for gametic *r*^2^ and zygotic *r*^2^ is as expected from equation 3.

**Figure 6.**
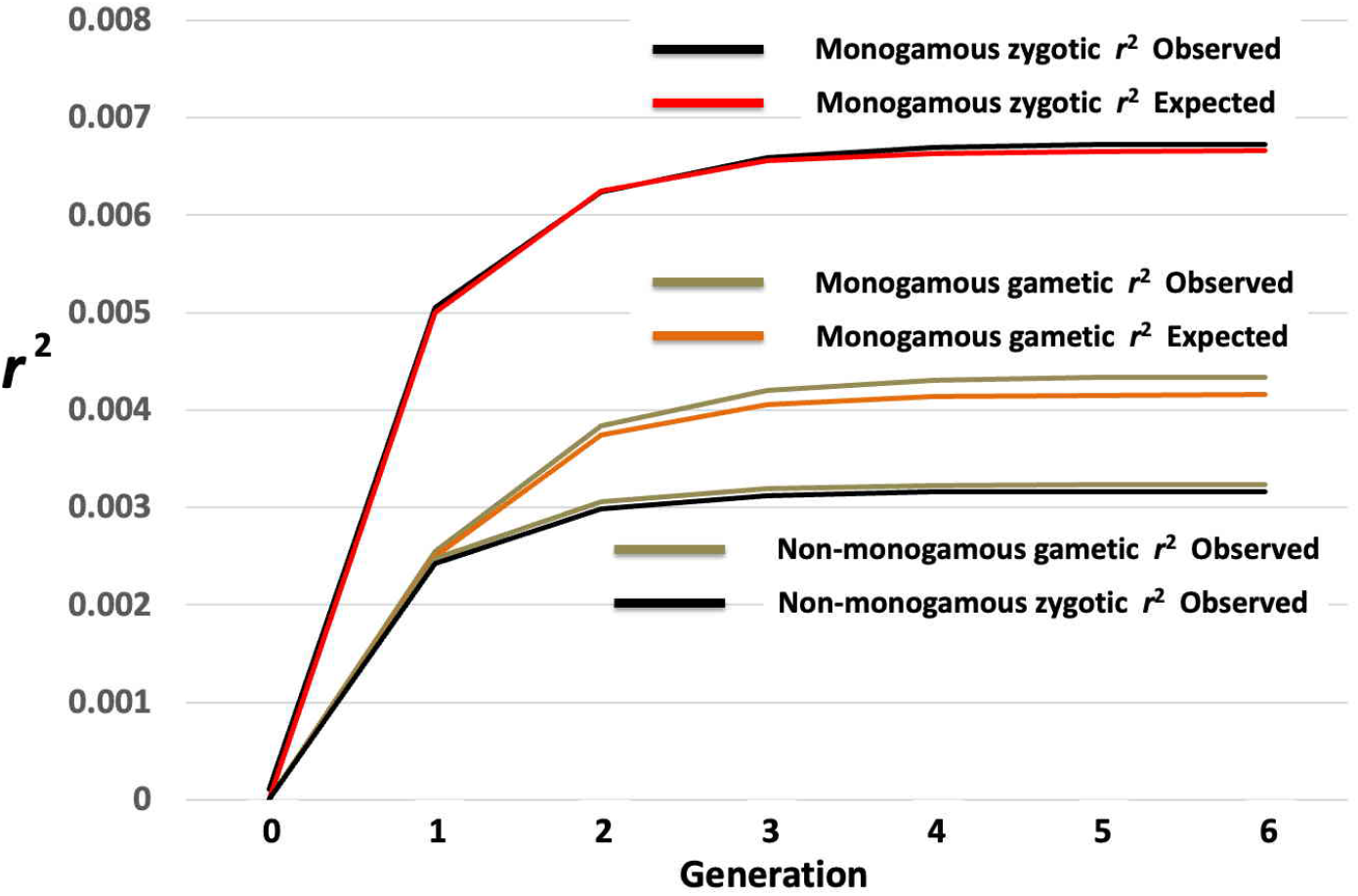
Observed and expected values for monogamous mating with gametic and zygotic *r*^2^ correlations.

The remaining graphs show the results of monogamous mating, including gametic and zygotic *r*^2^ and their expectations. They show that monogamous mating leads to a substantial increase in *r*^2^ values, as predicted by equations 16 and 18. The expectation for Generation 1 of Figure 6 uses equation 13, because equation 18 assumes an extra generation of mating that is not present in Generation 0.

## Discussion

The primary conclusion from this paper is that expectations for most LD formulae involving *r*^2^ can be derived using simple probability arguments. These show in a reasonably straightforward manner why the haploid and diploid mating systems lead to slightly different outcomes. Further, why random partner choice for each offspring versus lifetime partner choice (monogamy) leads to differences in LD expectation, and why this difference is particularly accentuated for zygotic *r*^2^ compared to gametic *r*^2^.

The designation “Linked Identity by Descent - LIBD” has previously been used [15] to describe the passage of haplotypes from a common ancestor, without recombination. The term has been used in the present paper, but with one qualification. The second panel of Figure 2 describes a situation where two haplotypes are passed down without recombination, but not haplotypes that themselves trace to a single haplotype. It accounts for the difference in outcome between the haploid and diploid model.

The key feature of Figure 2 is the coalescence to a single diploid individual. Coalescence is usually defined in terms of convergence to a single locus or haplotype [9], rather than to an individual. A term such as “Two-locus Diploid Coalescence” might therefore be more exact here. The LIBD descriptor, with its implication that linkage is necessarily involved, may seem less appropriate, particularly since the two loci of Figure 2 could be loci on separate chromosomes rather than linked to each other.

It is instructive to draw attention here to the term “Linkage Disequilibrium” that seemingly has become established through common usage. Some authors have objected to this term, largely because unlinked loci can also be in “disequilibrium” [1]. However, the major implications of LD come from the expectation of LD between SNPs and closely linked loci of interest. Multiple genome-wide association studies (GWAS) are based on this expectation [17]. Similarly, the use of genomic breeding values has revolutionised animal and plant breeding [11]. By comparison, correlations between unlinked and loosely linked loci are usually of secondary interest, although correlations between such loci may be specifically useful in conservation genetics to estimate effective population size [18].

We argue that the same situation pertains here. The LIBD calculation leads immediately to an accurate prediction for LD for closely linked loci (Figure 3). Only when linkage is sufficiently loose such that the term *c*^2^ becomes important, does the simulated *r*^2^ begin to diverge slightly from the LIBD-based expectation. On the other hand, the probabilistic definition of *r*^2^ as a LIBD measure offers an intuitive interpretation of the effective sample size (*Sr*^2^, where *S* is either the number of chromosomes or the number of individuals, depending on whether phased or unphased data are used) used in GWAS [13]: it corresponds to the expected number of sampled units carrying a shared, unbroken ancestral segment between the marker and the focal gene. Consequently, framing *r*^2^ through this genealogical lens clarifies how it isolates true, informative marker-gene associations from spurious signals driven merely by coincidence or identity-by-state (IBS).

## 6 Acknowledgement

Brian Charlesworth alerted us to Sewall Wright’s (1933) paper as the original source of the correlation definition of linkage disequilibrium.

## 7 Appendix

### A Calculations for two offspring sharing two parents (full sibs)

Looking initially at a single offspring individual, four offspring classes may be recognised, as shown with frequencies in Table S1. These classes are defined by Non-recombinant versus Recombinant for each of the two gametes, and each genotype as shown in the table is representative of four possible genotypes. Classes 2 and 3 are equivalent, but sometimes require separate treatment as seen below.

We now look at pairs of offspring. Since each offspring can be one of four classes, there are 5 *×* 4*/*2 = 10 possible pairs, each of which needs to be considered separately. Table S2 shows the calculations where both offspring are of Class 1. The end result is then shown as the first line in Table S3.

The first parent in Table S2 is arbitrarilily assigned the genotype 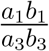. The calculation considers the probability that randomly selected haplotypes in Offspring 1 and Offspring 2 are identical. There are four such haplotype pairs for each pair of genotypes. For the first pair of individuals, both of genotype 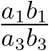 two haplotype pairs are identical, both *a*_1_*b*_1_ or both *a*_3_*b*_3_, giving a contribution of 2/4, to the LIBD probability.

**Table S1:** Four classes of offspring from parents 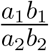 and 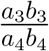. N and R refer to Non-recombinant and Recombinant.

| Class | Type | Genotype | Frequency |
| --- | --- | --- | --- |
| 1 | N,N | $\frac{a_1b_1}{a_3b_3}$ | $(1 - c)^2$ |
| 2 | N,R | $\frac{a_1b_1}{a_3b_4}$ | $c(1 - c)$ |
| 3 | R,N | $\frac{a_1b_2}{a_3b_3}$ | $c(1 - c)$ |
| 4 | R,R | $\frac{a_1b_2}{a_3b_4}$ | $c^2$ |

In two cases from these genotypes, the haplotypes are *a*_1_*b*_1_ and *a*_3_*b*_3_. LIBD is possible in this case if there was LIBD in the parents, whose LIBD probability is assigned as *L*.

Summing over the four classes, the overall contribution from the genotypes of Table S2 to *L*^*′*^ is:

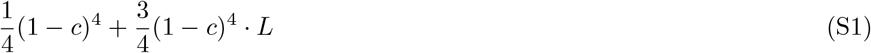

which is entered as the first line of Table S3.

Contributions from each possible pair of genotype classes to Table S3 can be calculated in the same way. One difference should be noted, specifically in cases where recombinant products occur in both offspring. Table S4, where both offspring are of Class 2, illustrates this. Genotype *a*_3_*b*_4_ here contributes to the ‘De novo LIBD’ in row 3. LIBD in this case occurs where the offspring come from the same parent, but not without recombination. This can be described as a case of ‘compensating crossovers’.

**Table S2:** Probabilities of offspring pairs where both are Class 1 (N, N)

| Offspring 1 | Offspring 2 | Frequency | De novo LIBD | Retained LIBD |
| --- | --- | --- | --- | --- |
| $\frac{a_1 b_1}{a_3 b_3}$ | $\frac{a_1 b_1}{a_3 b_3}$ | $\frac{(1-c)^4}{4}$ | $\frac{2}{4}$ | $\frac{2}{4}$ |
| $\frac{a_1 b_1}{a_3 b_3}$ | $\frac{a_1 b_1}{a_4 b_4}$ | $\frac{(1-c)^4}{4}$ | $\frac{1}{4}$ | $\frac{3}{4}$ |
| $\frac{a_1 b_1}{a_3 b_3}$ | $\frac{a_2 b_2}{a_3 b_3}$ | $\frac{(1-c)^4}{4}$ | $\frac{1}{4}$ | $\frac{3}{4}$ |
| $\frac{a_1 b_1}{a_3 b_3}$ | $\frac{a_2 b_2}{a_4 b_4}$ | $\frac{(1-c)^4}{4}$ | 0 | 1 |

**Table S3:**
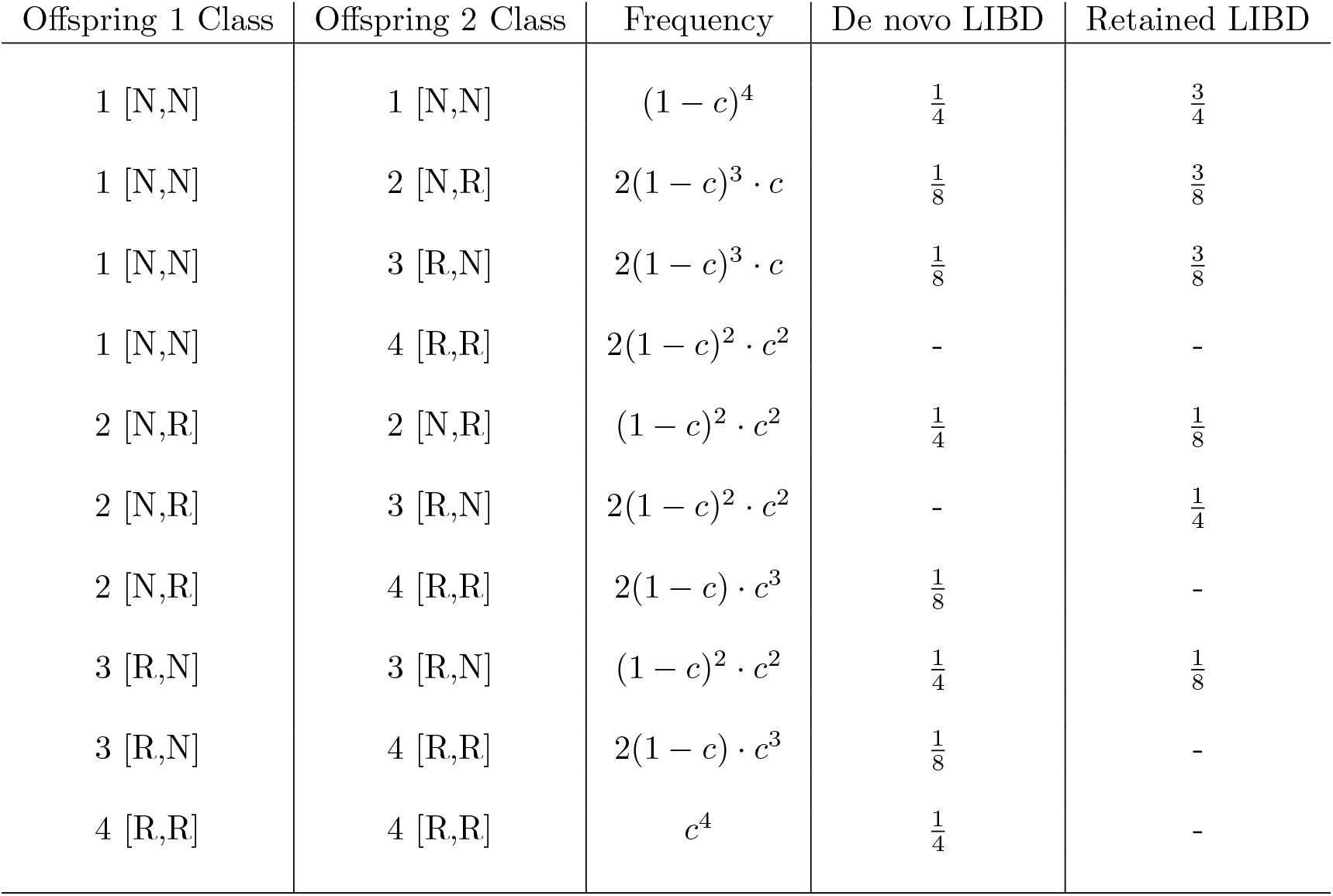
Summary of contributions to *L*^*′*^ from different genotype classes with two common parents.

| Offspring 1 Class | Offspring 2 Class | Frequency | De novo LIBD | Retained LIBD |
| --- | --- | --- | --- | --- |
| 1 [N,N] | 1 [N,N] | $(1-c)^4$ | $\frac{1}{4}$ | $\frac{3}{4}$ |
| 1 [N,N] | 2 [N,R] | $2(1-c)^3 \cdot c$ | $\frac{1}{8}$ | $\frac{3}{8}$ |
| 1 [N,N] | 3 [R,N] | $2(1-c)^3 \cdot c$ | $\frac{1}{8}$ | $\frac{3}{8}$ |
| 1 [N,N] | 4 [R,R] | $2(1-c)^2 \cdot c^2$ | - | - |
| 2 [N,R] | 2 [N,R] | $(1-c)^2 \cdot c^2$ | $\frac{1}{4}$ | $\frac{1}{8}$ |
| 2 [N,R] | 3 [R,N] | $2(1-c)^2 \cdot c^2$ | - | $\frac{1}{4}$ |
| 2 [N,R] | 4 [R,R] | $2(1-c) \cdot c^3$ | $\frac{1}{8}$ | - |
| 3 [R,N] | 3 [R,N] | $(1-c)^2 \cdot c^2$ | $\frac{1}{4}$ | $\frac{1}{8}$ |
| 3 [R,N] | 4 [R,R] | $2(1-c) \cdot c^3$ | $\frac{1}{8}$ | - |
| 4 [R,R] | 4 [R,R] | $c^4$ | $\frac{1}{4}$ | - |

**Table S4:** Probabilities of offspring pairs, both Class 2 (N, R)

| Offspring 1 | Offspring 2 | Frequency | De novo LIBD | Retained LIBD |
| --- | --- | --- | --- | --- |
| $\frac{a_1 b_1}{a_3 b_4}$ | $\frac{a_1 b_1}{a_3 b_4}$ | $\frac{c(1-c)^3}{2}$ | $\frac{2}{4}$ | 0 |
| $\frac{a_1 b_1}{a_3 b_4}$ | $\frac{a_1 b_1}{a_4 b_3}$ | $\frac{c(1-c)^3}{2}$ | $\frac{1}{4}$ | 0 |
| $\frac{a_1 b_1}{a_3 b_4}$ | $\frac{a_2 b_2}{a_3 b_4}$ | $\frac{c(1-c)^3}{2}$ | $\frac{1}{4}$ | $\frac{1}{4}$ |
| $\frac{a_1 b_1}{a_3 b_4}$ | $\frac{a_2 b_2}{a_4 b_3}$ | $\frac{c(1-c)^3}{2}$ | 0 | $\frac{1}{4}$ |

Summing all contributions to *L*^*′*^ from Table S3 gives

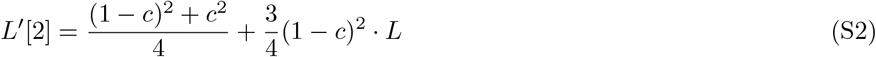

where the symbol *L*^*′*^[2] denotes the LIBD offspring probability when there are two common parents. It is noteworthy that all terms in *c*^3^ and *c*^4^ cancel to give this simple result, presumably attributable to the fact that *L* is ultimately a statistic referring to gametes rather than genotypes.

Equation S2 is entered as equation 10 in the main text.

### B Calculations for two offspring sharing one parent (half sibs)

The same calculations as summarised in Table S3 are shown in Table S5 for the single common parent case

**Table S5:** Summary of contributions to *L*^*′*^ from different genotype pairs with a single common parent.

| Offspring 1 | Offspring 2 | Frequency | De novo LIBD | Retained LIBD |
| --- | --- | --- | --- | --- |
| 1 [N,N] | 1 [N,N] | $(1 - c)^4$ | $\frac{1}{8}$ | $\frac{7}{8}$ |
| 1 [N,N] | 2 [N,R] | $2(1 - c)^3 \cdot c$ | - | $\frac{1}{2}$ |
| 1 [N,N] | 3 [R,N] | $2(1 - c)^3 \cdot c$ | $\frac{1}{8}$ | $\frac{3}{4}$ |
| 1 [N,N] | 4 [R,R] | $2(1 - c)^2 \cdot c^2$ | - | - |
| 2 [N,R] | 2 [N,R] | $(1 - c)^2 \cdot c^2$ | $\frac{1}{8}$ | $\frac{1}{4}$ |
| 2 [N,R] | 3 [R,N] | $2(1 - c)^2 \cdot c^2$ | - | $\frac{1}{4}$ |
| 2 [N,R] | 4 [R,R] | $2(1 - c) \cdot c^3$ | $\frac{1}{8}$ | - |
| 3 [R,N] | 3 [R,N] | $(1 - c)^2 \cdot c^2$ | $\frac{1}{8}$ | $\frac{1}{8}$ |
| 3 [R,N] | 4 [R,R] | $2(1 - c) \cdot c^3$ | - | - |
| 4 [R,R] | 4 [R,R] | $c^4$ | $\frac{1}{8}$ | - |

Summing contributions to *L*^*′*^ from Table S5 gives

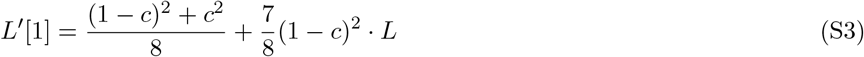

### C Innovation–preservation recursion for LIBD

Let *L*_*t*_ denote the probability that two sampled haplotypes at generation *t* exhibit the shared two-locus association state due to LIBD. This state may arise either through historical preservation of an existing two-locus association or through de novo generation during parental sampling.

The de novo contribution has two distinct pathways. Identical non-recombinant transmissions contribute

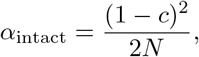

whereas identical recombinant transmissions contribute

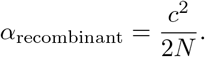

Thus, the total de novo association contribution due to LIBD is

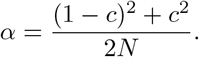

An existing historical association due to LIBD is retained when the two ancestral lineages do not coalesce to the same ancestral gamete and neither lineage recombines. Hence the historical preservation factor is

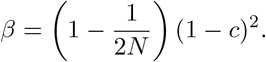

The dynamics of association due to LIBD is therefore

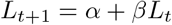

or, explicitly,

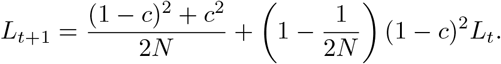

At equilibrium, *L*_*t*+1_ = *L*_*t*_ = *L*^*∗*^, giving

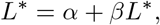

and hence

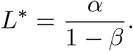

Substitution gives

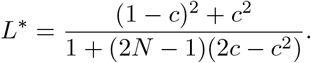

Thus, equilibrium association reflects a balance between de novo generation of shared two-locus states, *α*, and the loss of historically retained association, (1 *− β*)*L*_*t*_. These are equal when *L*_*t*_ = *L*^*∗*^.

### D Population simulation of a chromosome with recombination segments

a. The forward generation program ‘chrom sim’ simulates a single chromosome. An initial single-sex population contains N diploid individuals, each with two individually marked chromosomes. The population, with N=1000 for Figure 3, therefore starts with complete LD, *r*^2^ = 1. The chromosome is assumed to be of length 100cM. On average, therefore, there will be one recombination event per generation, at a random position on the chromosome. After *t* generations, each chromosome will contain on average *t* segments, each marked with an initial chromosome number (1 .. 2000). Note that these segments may be formed by recombination within regions of identity between the two chromosomes, so that they are not ‘junctions’ in the usually accepted version of the term. Each segment needs to be classified for each chromosome. A diploid population of 2,000 chromosomes simulated for 2,000 generations, as in Figure 3, requires classification of around 4,000,000 segments. The chrom sim program primarily simulates runs of segment identity. Calculation of *r*^2^ for each value of *Nc* involves selecting a random chromosome position for the first segment (*x*) and a second segment (*y*) at the required distance given by *c*. For each chromosome in the population *x* and *y* values are the segment markers (1 .. 2000) for the chosen segment of that particular chromosome. The value of *r*^2^ is then calculated, and the procedure repeated 1,000 times to give an average *r*^2^. Each population is simulated 100 times for an overall average *r*^2^. Note that the correlation here is between multiple values of *x* and *y* rather than just two values as considered in Tables 1 and 2.
b. The ‘2loc sim’ program is a simple 2-locus 2-allele simulation, starting from linkage equilibrium. The program allows both haploid and diploid simulations. Diploid simulations can be either single sex or two sexes. Within the latter, monogamous mating can be specified. At each generation, both gametic and zygotic *r*^2^ values are calculated,

**Table 1:**
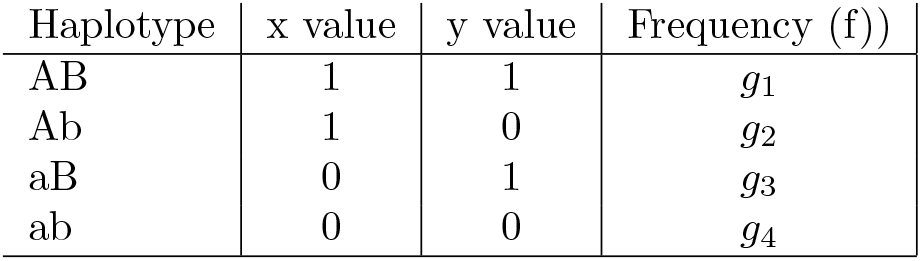
Gametic covariance table.

**Table 2:**
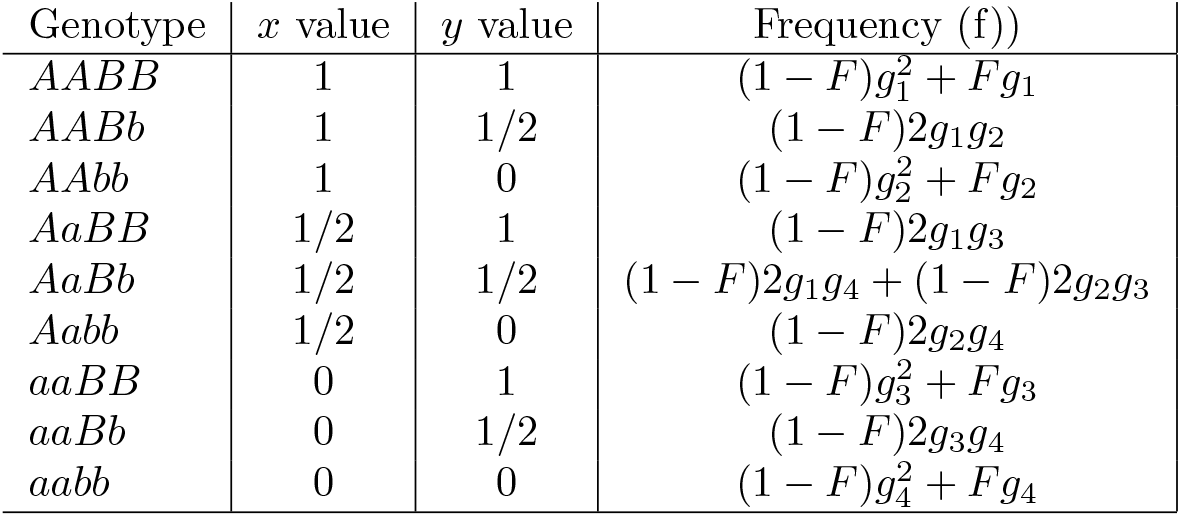
Zygotic covariance table.

**Table 3:** Parental classes (2, 1 or 0) and LIBD probabilities.

| Common parent(s) | Frequency | L' | De novo LIBD | Retained LIBD |
| --- | --- | --- | --- | --- |
| Both | $\frac{1}{N_f} \frac{1}{N_m}$ | $L'[2]$ | $\frac{(1-c)^2+c^2}{4}$ | $\frac{3}{4}(1-c)^2 \cdot L$ |
| Female | $\frac{1}{N_f}(1 - \frac{1}{N_m})$ | $L'[1]$ | $\frac{(1-c)^2+c^2}{8}$ | $\frac{7}{8}(1-c)^2 \cdot L$ |
| Male | $(1 - \frac{1}{N_f})\frac{1}{N_m}$ | $L'[1]$ | $\frac{(1-c)^2+c^2}{8}$ | $\frac{7}{8}(1-c)^2 \cdot L$ |
| Neither | $(1 - \frac{1}{N_f})(1 - \frac{1}{N_m})$ | $L'[0]$ | - | $(1-c)^2 \cdot L$ |

**Table 4:** Parental classes and LIBD probabilities for monogamous mating.

| Common parent(s) | Frequency | L' | De novo LIBD | Retained LIBD |
| --- | --- | --- | --- | --- |
| Both | $\frac{2}{N_e}$ | $L'[2]$ | $\frac{(1-c)^2+c^2}{4}$ | $\frac{3}{4}(1-c)^2 \cdot L$ |
| Neither | $1 - \frac{2}{N_e}$ | $L'[0]$ | - | $(1-c)^2 \cdot L$ |

This program requires a choice of starting allele frequency. Figure 4 assumes an initial starting frequency of 0.5 for each of two alleles, assigned independently at *A* and *B* loci. An advantage of the ‘chrom sim’ program is that it makes no assumption of initial frequency, although it requires an initial starting value of *r*^2^ = 1.

The importance of the frequency assumption in 2loc sim has been tested by varying the allele frequency to 0.4, 0.3, 0.2 and 0.1, Above approximately 0.25 there is little difference in the *r*^2^ output. Below this value, the *r*^2^ value is reduced, although not substantially. But below 0.25, fixation starts to occur, thereby complicating expectations (see equation 6 and associated text).

Individual runs vary widely. The results of figure 4 are averaged over 10^8^ replicate population simulations to reduce variability as much as possible.

